# Towards automated control of embryonic stem cell pluripotency

**DOI:** 10.1101/685297

**Authors:** Mahmoud Khazim, Lorena Postiglione, Elisa Pedone, Dan L. Rocca, Carine Zahra, Lucia Marucci

## Abstract

Mouse embryonic stem cells (mESCs) have been shown to exist in three distinct pluripotent states (ground, naïve and primed pluripotent states), depending on culture conditions. External feedback control strategies have been, so far, mainly used to automatically regulate gene expression in bacteria and yeast. Here, we exploit a microfluidics/microscopy platform and segmentation and external feedback control algorithms for the automatic regulation of pluripotency phenotypes in mESCs. We show feasibility of automatically controlling, in living mESCs, levels of an endogenous pluripotency gene, Rex1, through a fluorescent reporter, used as control output, and drugs commonly used to modulate pluripotency (i.e. MEK kinase and Gsk3β inhibitors) as control inputs. Our results will ultimately aid in the derivation of superior protocols for pluripotency maintenance and differentiation of mouse and human stem cells.

## 1. INTRODUCTION

Pluripotent stem cells are characterised by the self-renewal property (i.e. they can indefinitely produce daughter cells which maintain the progenitor cell properties), and by the ability to differentiate into all three germ layers (mesoderm, endoderm, and ectoderm). Pluripotency can be sustained indefinitely in *in vitro* cultures, if adequate cultures are established.

Isogenic mouse embryonic stem cells (mESCs) can exist in varying pluripotency states depending on culture conditions (Hackett et al., 2014). mESCs maintained in serum/LIF media, containing serum and the leukemia inhibitory factor (LIF), are in a naïve pluripotency state and display heterogeneous expression of core pluripotency genes including Nanog, Rex1, Esrrb and β-catenin (Hatano et al., 2005, Chambers et al., 2007, Toyooka et al., 2008, Hayashi et al., 2008, Van Den Berg et al., 2008, Marucci et al., 2014). The addition of the two small molecule inhibitors PD0325901 (PD, MEK kinase inhibitor) and Chir99021 (Chiron, a GSK3β inhibitor) in the so-called 2i/LIF media (which is serum free, but still contains LIF) has been shown to push mESCs into a ground state of pluripotency (Fig. 1), producing a significant upregulation in the expression of key pluripotency genes (Kolodziejczyk et al., 2015, Wray et al., 2011). The naïve and ground states exhibit distinct transcriptional profiles (Ghimire et al., 2018, Marks et al., 2012) and are thought to influence lineage specification and differentiation priming (Kalkan et al., 2014, Loh et al., 2011), and to alter cellular response to differentiation cues (Abranches et al., 2014). Still, the overall homogeneity of pluripotency genes in 2i/LIF cultures has been challenged (Morgani et al., 2013, Abranches et al., 2014, Godwin et al., 2017), and long-term mESC cultures in 2i/LIF have been recently shown to present genomic instability (Choi et al., 2017).

**Fig. 1.**
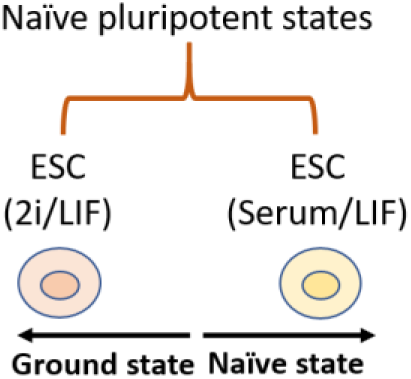
Naïve pluripotent states: mESCs derived in conventional culture conditions, which include the leukemia inhibitory factor (LIF) and fetal bovine serum, are representatives of the naïve state of pluripotency, with fluctuating expression of pluripotency genes. mESCs cultured in a medium supplemented with 2i (two inhibitors of MEK and GSK3β) and LIF represent the ground state of pluripotency.

External feedback control strategies have been recently applied to biological system to automatically tune gene expression in living cells (Del Vecchio et al., 2016, Danino et al., 2010, Milias-Argeitis et al., 2011, Uhlendorf et al., 2012, Menolascina et al., 2014, Fiore et al., 2016, Milias-Argeitis et al., 2016, Toettcher et al., 2011, Fracassi et al., 2016).

Our lab has experience in employing feedback control paradigms to control mammalian cell behaviour (Pedone et al., 2018, Postiglione et al., 2018). By using a microfluidics/microscopy platform to precisely manipulate culture conditions while measuring gene expression, we have been able to control various mammalian cell lines towards desired behaviours. This strategy is implemented by measuring, over a time-lapse experiment, the system output (fluorescence, as a proxy for gene expression), which is compared to a control reference (desired fluorescence). This difference is termed the control error and is used by the control algorithm to determine the control input. The platform is able to manipulate, over the time-lapse, the cellular microenvironment through actuators, which provide to cells, cultured in a PDMS microfluidics device (Kolnik et al., 2012), time- and magnitude-varying stimuli as control inputs (Fig. 2).

**Fig. 2.**
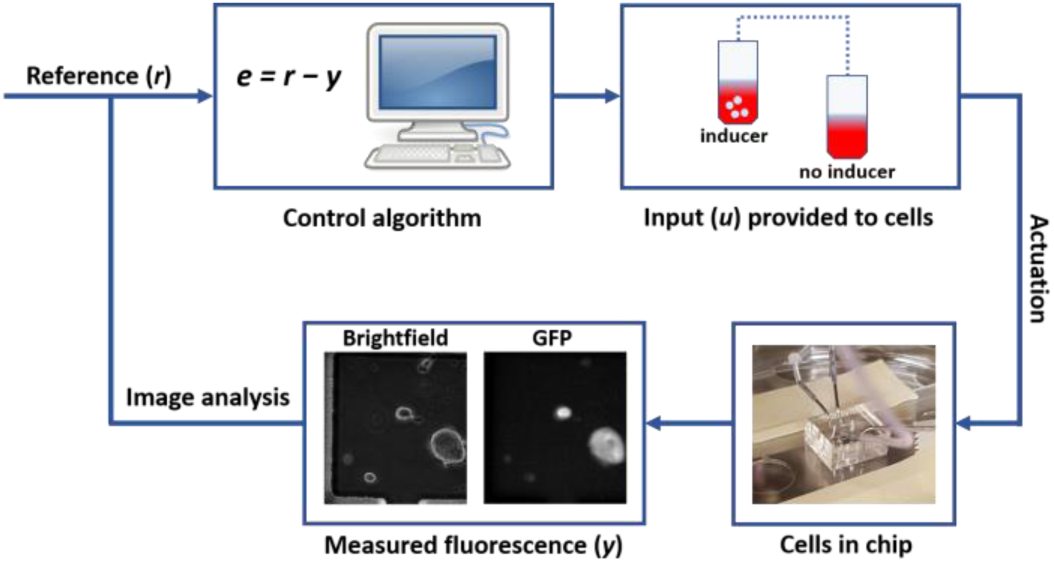
The microfluidics/microscopy platform for external feedback control. The platform consists of: i) the microfluidic device with mESCs growing within; ii) an inverted widefield microscope that takes phase contrast and fluorescent images; iii) a computer implementing segmentation and control algorithms, which measure the fluorescence (y), and calculate the control error (e) and control input (u) to direct the system towards a pre-set reference fluorescence (r); iv) an actuation system providing cells media with/without control input.

Here, we show preliminary results towards application of this technology for the direct engineering of cell fate in mESCs. We used a mESC line containing a Rex1GFPd2 reporter wherein one allele of the Rex1 coding sequence is replaced with the coding sequence of a destabilised green fluorescent protein (GFP), a fluorescent reporter with relatively short half-life of 2-hours (Fig. 3a). The Rex1 gene is a marker for undifferentiated stem cells and is regulated by Nanog, Sox2 and Oct4 transcriptional regulators of pluripotency (Kashyap et al., 2009). Rex1 expression has been noted to fluctuate, with a multimodal distribution observed in conventional serum/LIF culture conditions (Hatano et al., 2005). Rex1 fluctuations and heterogeneity are reduced by the addition of the two inhibitors Chiron and PD (Wray et al., 2010, Kalkan et al., 2017).

**Fig. 3.**
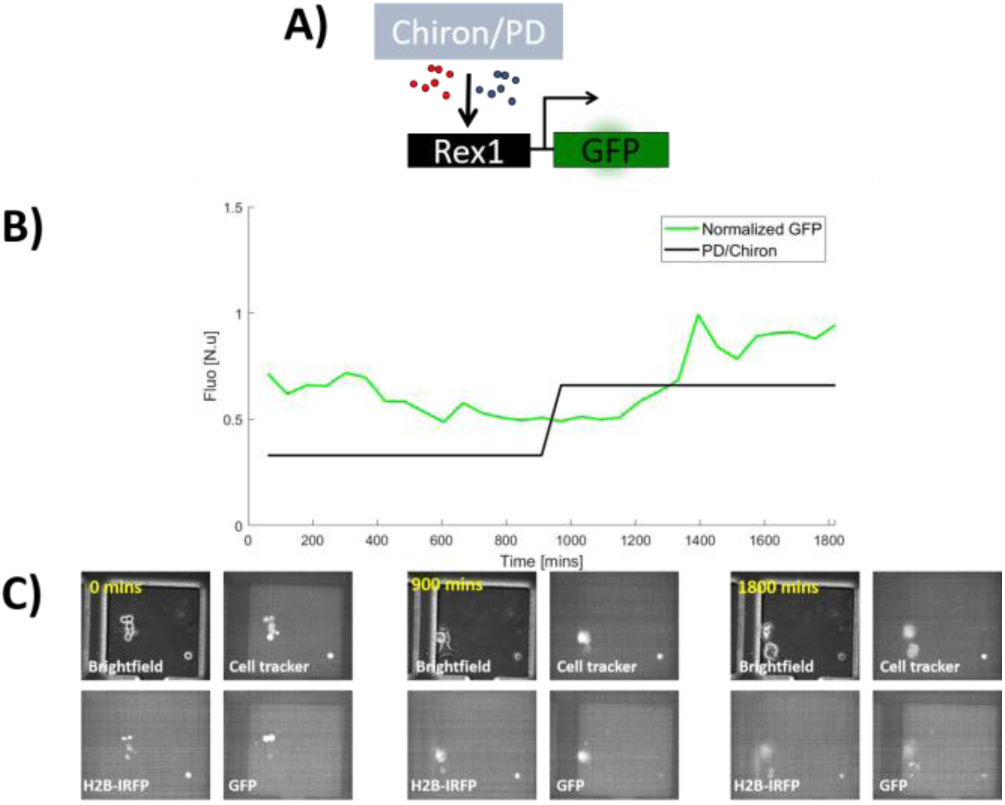
Step-up open loop experiment. A) mESCs carrying a mono-allelic GFP knock-in at the Rex1 (pluripotency marker) locus used in the experiment. B) Rex1-GFPd2 mESCs, carrying a H2B-iRFP nuclear marker, were perfused for 16 hours in Serum/LIF media and cell tracker dye. The experiment was initialised with a switch to media containing 0.33µM of PD and 1µM Chiron for 15 hours and then 0.66µM of PD and 2µM Chiron for the following 15 hours. Increasing concentrations of PD/Chiron induced an increase in GFP expression during transition towards ground state. C) Representative brightfield, Cell tracker (Blue), H2B-iRFP and GFP images taken at 0, 900 and 1800 mins.

We show feasibility of automatic controlling Rex1 expression in an isogenic population of mESCs; these preliminary results suggest that cell fate can be automatically and precisely tuned by exploiting feedback strategies. Also, we present the design of a novel microfluidic device for culture, imaging and control of human pluripotent cells.

Our results should help in the production of novel protocols for increased efficiency of pluripotency maintenance and differentiation in a cell response-informed way. The proposed methodologies should be easy to adapt to other cell lines and processes (e.g. cancer stem cells and differentiation, respectively).

## 2. METHODS

### 2.1 Cell culture and transfection

The mESC line used in this study is the Rex1-GFP reporter line (Wray et al., 2011). Cells were cultured on gelatinised tissue culture dishes at 37°C in a 5% CO_2_ humidified incubator in Dulbecco’s modified Eagle’s medium (DMEM) supplemented with 15% fetal bovine serum (Sigma), non-essential amino acids, L-glutamine, sodium pyruvate, Penicillin-Streptomycin, 2-mercaptoethanol and 10ng/ml mLIF (Peprotech; 250-02).

For the open-loop experiment shown in Fig. 3, cells were perfused overnight in serum/LIF; when the time-lapse was started, mESCs were given, in 15-hour steps, increasing concentrations of inducers (0.33µM PD (Stratech; S1036-SEL) and 1µM Chiron (Stratech; S1263-SEL) and then 0.66µM PD and 2µM Chiron, respectively). For the Relay control experiment in Fig. 4, cells in the microfluidic chip were perfused overnight with media supplemented with 1µM PD and 3µM Chiron.

**Fig. 4.**
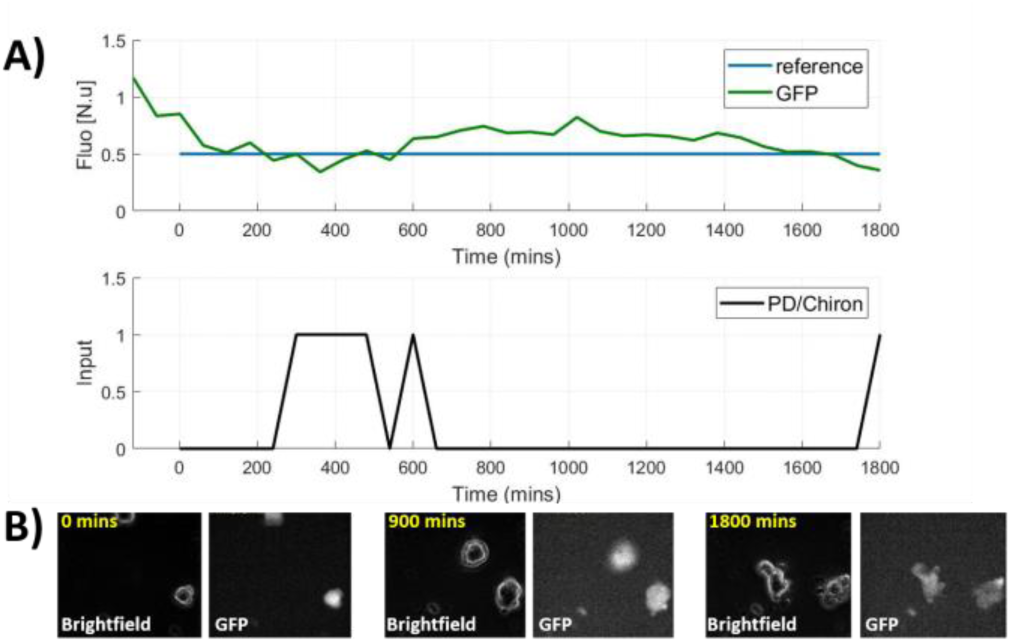
Set-point Relay control experiment. A) mESCs were cultured overnight with PD/Chiron; after control experiment initialisation, PD/Chiron were provided by actuators given the current GFP fluorescence, using a Relay algorithm (see Methods). B) Representative brightfield and GFP fluorescence images at 0, 900 and 1800 minutes.

### 2.2 Transfection with nuclear marker and single cell sorting

mESCs in microfluidic chip show a degree of morphological heterogeneity. To improve segmentation, mESCs were transfected with a plasmid to express a transgene to encode Histone 2B (H2B) tagged with infra-red fluorescent protein (iRFP), as well as hygromycin resistance cassette, using 2µg of plasmid DNA and Polyfect transfection reagent (Qiagen). Positive selection was performed by adding 25µg/ml of hygromycin B (Sigma) to culture media. Furthermore, clonal FACS sorting was performed to produce cells which homogeneously expressed the iRFP-tagged H2B protein.

### 2.3 Microfluidic chip fabrication

The microfluidic device for mammalian cells was designed and characterised by Kolnik and colleagues. (Kolnik et al., 2012). A master mould of the device was purchased (MicruX technologies), and replica were produced by PDMS moulding. Briefly, PDMS monomer and curing agent (Sylgard 184, Dow Corning, Ellsworth Adhesives) were mixed together at a 1:10 ratio, as per manufacturer’s instructions. This mixture was poured over the master and degassed until all bubbles were removed. The mixture was then cured at 80°C for an hour. The cured PDMS stamp was peeled off the master and dry autoclaved at 121°C for 30 mins. A scalpel was used to cut the microfluidic devices from the PDMS and a 0.75mm biopsy punch used to create holes which serve as the ports of the device. After punching, chips were sonicated in isopropyl alcohol for 10 mins followed by sonication in dH2O for 10 mins. These devices were then bonded to thin glass coverslips by exposing both device and coverslip to oxygen plasma for 1 min and contact bonded.

### 2.4 Loading of cells and fluorescent time-lapse imaging

Cells were loaded by vacuum pump as described in (Kolnik et al., 2012), and allowed to attach overnight with constant perfusion of fresh media in a humidified incubator. The device was then secured onto the microscope stage within an incubation chamber (incubator i8; Leica microsystems) maintaining the device at 37°C in a 5% CO_2_; 60ml syringes were connected via tubing to inlet and waste ports, with hydrostatic pressure used to maintain the proper flow of medium. The syringes, attached to the inlet ports, were secured onto linear motorised actuators (L C Automation Ltd.) with induction and non-induction media in respective axes. Time-lapse imaging to acquire phase and fluorescent images was obtained using Leica LASX live cell imaging workstation on a DMi8 inverted fluorescent microscope with an Andor iXON 897 Ultra digital camera using a x40 objective (PlanFluor DLL 40x Ph2 Nikon). A motorised stage and adaptive focus control were used to choose fields to image. Flow properties were monitored by the addition of 1µM Sulforhodamine 101 (Sigma) in medium containing the inducer molecules. The set-up for the experiment in Fig. 4 consisted of 3-channel acquisition (Phase contrast, Green and Red fluorescence) every 60 mins; in the experiment in Fig. 3, which includes both the nuclear H2B-iRFP tag and cell tracker dye (Invitrogen; C12881), Blue and Infra-red fluorescent filter cubes were also used.

### 2.5 Segmentation and control algorithms

Briefly, the segmentation algorithm (programmed in Matlab -Mathworks Matlab R2018a-) utilises image-processing functions (i.e. threshold definition to generate a binary image for the cell/colony edge, and dilation and filling operators) to create a mask. This mask is applied to the fluorescent images. In the experiment in Fig. 3, where mESCs carrying a H2B-iRFP tag were imaged in presence of the tracking dye, the mask was calculated using the cell tracker dye; the average GFP intensity was then normalised to iRFP intensity. In the experiment in Fig. 4, the mask was generated from phase contrast images and the average GFP intensity was normalised to the mask area to account for varying cell number over the time-lapse. In both experimental set-ups, to account for possible fluctuations in the background, the fluorescent intensity was calculated also on a portion of the field free of cells; this was then subtracted from the quantified GFP intensity. In the closed-loop experiment in Fig. 4, the max GFP value, normalised to 1, was calculated by averaging GFP intensity over 2 hours of initialisation (i.e. mESCs, pre-treated for 16 hours with 1µM PD+3µM Chiron, were kept in the platform for 2 hours in presence of both inducers prior to closed loop experiment start).

To maintain the system at the desired fluorescence in the closed loop experiment (Fig. 4), a Relay control strategy was employed. The control error *e(t)* was calculated at each sampling time (60 minutes) using the following equation:

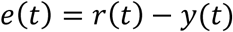

which calculates the difference between the desired reference fluorescence (*r*) and the system output (*y*) at time (*t*). The control law (*u*) can be expressed as follows:

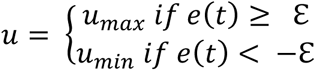

where the hysteresis ε is equal to 5% of reference value. In the experiment in Fig. 4, *u*_*max*_ corresponds to 1µM PD+3µM Chiron enriched media, while *u*_*min*_ corresponds to plain media.

### 2.6 Novel microfluidic design simulations

Modifications of the original microfluidic device design (Kolnik et al., 2012) were performed using AutoCAD (Autodesk) with the main design feature being increased inlet and chamber sizes so as to allow human pluripotent stem cell clusters to enter the chambers. Finite element analysis was performed in COMSOL Multiphysics (Version 5.3.0) to calculate the mean fluid velocity in the new designs.

## 3. RESULTS

### 3.1 Open loop testing of the system response

Firstly, we tested in our microfluidics/microscopy platform the response of the fluorescence GFP reporter, tagging the endogenous pluripotency gene Rex1 in the aforementioned mESC line (Fig. 3a), to pluripotency inducers. Cells were perfused for 16 hours overnight with serum/LIF and the cell permeable blue tracker. Once time-lapse imaging and fluorescence measurements were initiated, increasing concentrations of PD/Chiron were delivered to cells (Fig. 3b, c). Cells showed a good response to varying inducer concentrations, indicating feasibility of mESC culture, imaging and actuation in our platform.

### 3.2 Relay control of an endogenous pluripotency gene in mESCs

We then attempted a feedback control experiment, using a Relay control algorithm with a set-point reference: mESCs were controlled to reach and maintain 50% of the initial fluorescence, with a sampling time of 60 minutes. Cells were perfused for 16 hours overnight with induction factors PD and Chiron, in basal serum/LIF media, to reach the maximum steady state activation of Rex1GFP. Once the control experiment was initialised, as expected, PD and Chiron were initially removed until the desired reference fluorescence was achieved; pulses of PD and Chiron were then delivered to cells following a Relay strategy (Fig. 4a). mESCs showed good response to drugs and the feedback strategy, although simple, was effective to automatically regulate Rex1 expression (Fig. 4a, b).

### 3.3 Development of a microfluidic device for human pluripotent stem cells

PDMS-based microfluidic devices have been designed for human pluripotent stem cell (hPSC) cultures (see (Zhang et al., 2017) for a review). Nevertheless, of the previously proposed devices (e.g. (Gómez-Sjöberg et al., 2007)), those enabling mixing of different inputs are often not adequate to generate complex and time-varying input profiles, which is desirable when the aim is to apply external feedback control. Therefore, we tested the microfluidic device we used for mESC (Kolnik et al., 2012), which enables precise on-chip mixing of two media through a dial-a-wave generator and fast media diffusion to chamber, to culture hPSCs. hPSCs are routinely grown as colonies in dish cultures. As such, attempts to use the original microfluidic device (Kolnik et al., 2012) with these cells required single cell dissociation due to inlet size restrictions, resulting in a reduction in cell viability (data not shown). Thus, design modifications were implemented to increase both inlet and chamber size and to allow aggregates to enter the chambers; we also designed modifications to the main perfusion channel size. AutoCAD was employed to modify the device, with several design options interrogated (Fig. 5a). The new designs were informed by COMSOL simulations to compare flow properties with those of the original device (Fig. 5a).

**Fig. 5.**
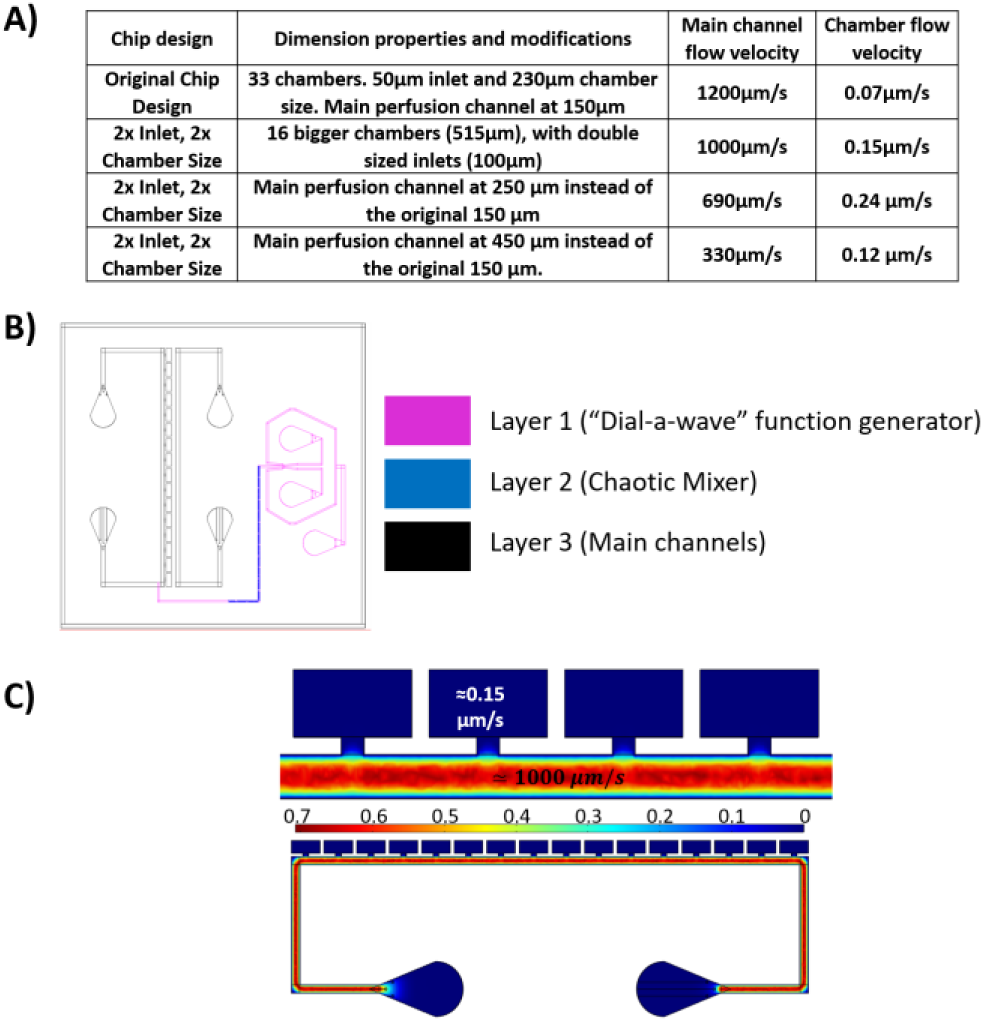
Design of a modified device modified for hPSC culture. A) COMSOL finite elements simulations showing main perfusion channel and chamber flow velocities for original device and modified designs. B) AutoCAD design for the 2x Inlet size, 2x Chamber size design modification. The design consists of three layers of different heights. All other features retain their unmodified dimensions from (Kolnik et al., 2012). C) Velocity flow rate in the 2x Inlet size, 2x Chamber. The flow is 1000μm/s in the main perfusion channel and 0.15μm/s in the chamber, with a 100μm wide inlet. Heat chart represents flow velocity magnitude (mm/s).

For preliminary operational characterisation, the simple double inlet, double chamber size design was taken forward to mask printing for fabrication. This design (Fig. 5b) showed a slight decrease in main perfusion flow velocity (1200µm/s to 1000µm/s), as well as an increase in chamber flow (from 0.07µm/s to 0.15µm/s), as compared to the original device design (Fig. 5c). These changes in flow characteristic are hypothesised to do not affect cell viability from increased shear stress and/or the ability of the device platform to generate continuous waveforms. For mask alignment of the three layers, an alignment strategy for multi-layered soft-lithography based on (Heymann et al., 2014) was included in the AutoCAD design, which includes corresponding crosses and squares and a Vernier scale.

## 4. DISCUSSION

Ground state pluripotency of mESCs in 2i/LIF cultures has been linked to cells expressing homogeneously higher levels of key pluripotency genes with a decrease in expression of differentiation markers as compared to conventional serum/LIF culture (Trott et al., 2013). ESCs are thought of as transcriptionally hyperactive (Efroni et al., 2008), and promiscuous transcription might be related to lineage priming (Loh et al., 2011). These states may thus bias the efficiency of lineage commitment. Our vision is that automatic control will help in the development of standardised cultures, thanks to the ability of gaining full control of key pluripotency and differentiation regulators.

Here, we described the use of feedback control theory principles for the automatic regulation of *in vitro* stem cell dynamics. Preliminary results show feasibility of our strategy, using a cellular paradigm for the control of naïve stem cell states. We are currently performing further experiments in mESCs, employing different cell lines and control algorithms, and developing new strategies for the implementation of feedback control in hPSCs.

## 5. ACKNOWLEDGMENTS

This work was funded by Medical Research Council grant MR/N021444/1 to LM, by the Engineering and Physical Sciences Research Council grant EP/R041695/1 to LM, and by BrisSynBio, a BBSRC/EPSRC Synthetic Biology Research Centre (BB/L01386X/1) to LM.

